# Radiolabeling of PSMA-617 with ^89^Zr: A Novel Use of DMSO for Radiochemical Yield Enhancement and Preliminary Small-Animal PET Results

**DOI:** 10.1101/2021.06.28.450175

**Authors:** Ryota Imura, Atsuko Nakanishi Ozeki, Nanako Shida, Mika Kobayashi, Hiroyuki Ida, Youichiro Wada, Nobuyoshi Akimitsu, Yoshitaka Kumakura

## Abstract

**Introduction:** Prostate-specific membrane antigen (PSMA)-targeted ligands, including PSMA-617, have been developed for theranostics of prostate cancer. ^68^Ga-PSMA-617 is the de facto standard of PSMA Positron Emission Tomography (PET) for imaging in prostate cancer patients prior to radioligand therapy (RLT) with ^177^Lu-PSMA-617. The dose-limiting toxicity for PSMA-RLT is damage to the kidney. PET scans using ^68^Ga-PSMA-617 have to be performed within a few hours of injection due to its short half-life (68 min). However, the presence of radioactivity in urine at the PET imaging timepoint hampers the dose optimization of ^177^Lu (half-life 6.6 d)-labeled PSMA-617. Thus, the long-lived positron emitter ^89^Zr (half-life 3.3 d) is suited for optimizing the doses of ^177^Lu-PSMA-617 because PET scans can be performed after excretion of radioactive urine. Although ^89^Zr has great potential for PET imaging, its inadequate incorporation into 1,4,7,10-tetraazacyclododecane-1,4,7,10-tetraacetic acid (DOTA), limits its applications. Here, we report the radiolabeling of PSMA-617 with ^89^Zr and preliminary PET imaging studies using ^89^Zr-PSMA-617.

**Methods:** DMSO and HEPES buffer were used to label PSMA-617 with ^89^Zr. The dissociation constant (*K*_d_) of ^89^Zr-PSMA-617 was determined using a cell-binding assay. Delayed-PET scans using ^89^Zr-PSMA-617 were performed at 24 h (N = 5).

**Results:** ^89^Zr-PSMA-617 was prepared with a radiochemical yield of 70 ± 9%. The *K*_d_ value was 6.8 nM. In PET imaging, standardized uptake value (SUV) was highest in LNCaP tumors (SUV_max_ = 0.98 ± 0.32), whereas it was low in kidney (SUV_max_ = 0.18 ± 0.7).

**Conclusion:** The preparation of ^89^Zr-PSMA-617 was achieved by using the DMSO and HEPES buffer. ^89^Zr-PSMA-617 visualize the PSMA positive LNCaP tumors without accumulation in bladder.

**Advances in knowledge and implications for patient care:** The use of ^89^Zr-PSMA-617 to predict the radiation doses in normal tissues lead to safe and effective RLT with ^177^Lu-PSMA-617.

## 1 Introduction

Prostate-specific membrane antigen (PSMA) is a transmembrane protein of 750 amino acids that is expressed in the prostate; some modest expression of PSMA is seen in normal tissues including the kidneys, salivary glands and lacrimal glands [1,2]. PSMA is overexpressed in prostate malignancy [3] and its expression is highest in metastatic castration-resistant prostate cancers (mCRPCs) that have a high Gleason score, which is indicative of an aggressive tumor. Thus, ligands that bind to PSMA-expressing cancer cells allow the targeted delivery of diagnostic and therapeutic agents [4–6]. One such small molecular ligand, PSMA-617, has potential utility in providing PSMA-targeted radioligand therapy (RLT) to patients with mCRPC [2,7]. PSMA-617 contains 1,4,7,10-tetraazacyclododecane-1,4,7,10-tetraacetic acid (DOTA), as a metal chelating moiety. PSMA-617 is most frequently conjugated to ^68^Ga and ^177^Lu as positron and beta ray-emitting radionuclides, respectively [2,8]. Thus, ^68^Ga-PSMA-617 is a de facto standard in PSMA PET for mCRPC, before RLT with ^177^Lu-PSMA-617.

Radionuclides with short half-lives are preferred for PET imaging because they expose patients to low radiation dose. However, the use of a positron emitter with a long half-life would provide advantages in terms of the ability to predict the radiation doses of normal tissues and organs at later time point. It is advantageous to compare them to the doses that are required to induce toxicity in mCRPC during PSMA-RLT. The dose-limiting toxicity of PSMA-RLT is set by the effects on the kidneys, because proximal renal tubular cells express PSMA. PET scans using ^68^Ga-PSMA-617 should be performed within a few hours from injection due to the short half-life of ^68^Ga (68 min). However, radioactivity persists in the urine at this time, which prevents dose optimization of ^177^Lu-labeled PSMA-617 (half-life 6.6 d). We hypothesized that the long-lived positron emitter ^89^Zr (half-life 3.3 d) would be better than ^68^Ga-PSMA-617 for optimizing the doses of ^177^Lu-PSMA-617, because PET scans could be performed after excretion of the radioactive urine. However, despite this potential advantage, the synthesis of ^89^Zr-PSMA-617 has not yet been achieved to the best of our knowledge.

Beyond the facilitation of dose optimization, the used of ^89^Zr offers additional advantages. First, its long half-life is ideal for delayed-PET scans. The tumor contrast against background is improved over time due to the clearing of non-specific radioligand distribution. Thus, PET scans at later time points from the injection (delayed-PET scans) yield high-contrast PET images. Indeed, high-contrast PET images were obtained by scanning patients at least 2–3 h after injection of ^68^Ga-PSMA-617 [2]. This delay can be extended up to 24 h or later if ^89^Zr-PSMA-617 is used. For this reason, higher contrast PET images are expected when ^89^Zr-PSMA-617 is used. Second, the relatively low positron energy of ^89^Zr (*E*_β+_: 396 keV, the mean positron range in water: 1.23 mm) improves spatial resolution compared to ^68^Ga (*E*_β+_: 836 keV, mean positron range in water: 3.48 mm) [9]. Third, a ready-to-use ^89^Zr-PSMA-617 formulation could be delivered from a central radiochemical facility; in contrast, ^68^Ga-labeled PSMA regents must be prepared under the radiochemist’s quality control in each hospital. Therefore, ^89^Zr-PSMA-617 may enable PSMA-targeted PET imaging even in hospitals that cannot afford to prepare ^68^Ga-labeled PSMA regents.

In this study, we hypothesized that ^89^Zr-PSMA-617 would be suited for optimizing the doses of ^177^Lu-PSMA-617. In what we believe to be the first report of its kind, we describe the radiolabeling of PSMA-617 with ^89^Zr in a mixture of aqueous buffer and organic solvent. In addition, we report a preliminary PET imaging study using ^89^Zr-PSMA-617.

## 2 Materials and methods

### 2.1 Production and purification of ^89^Zr

The ^89^Zr was produced from ^89^Y (natural abundance 100%) target via (p,n) reactions by a cyclotron (Cyclone 18, IBA). The radionuclide purity of irradiated ^89^Y targets was analyzed using a high-purity Ge detector (GMX15, Seiko EG&G). The produced ^89^Zr was purified by modified procedures described by Holland *et al.* [10]. Irradiated ^89^Y targets were dissolved in 2 mol/L HCl and loaded into a hydroxamate resin column. The column was washed with 10 mL of 2 mol/L HCl and 10 mL of water before elution of ^89^Zr with 3 mL of 1 mol/L oxalic acid. The eluate was loaded into Sep-Pak Accell Plus QMA Plus Light Cartridge (Waters). The cartridge was washed sequentially with 50 mL of water, 1 mL of 0.025 mol/L HCl, 1 mL of 0.05 mol/L HCl, and 0.3 mL of 0.1 mol/L HCl. Then, the ^89^Zr chloride solution was recovered by addition of 0.6 mL of 0.1 mol/L HCl.

The oxalic acid remaining in the purified ^89^Zr chloride solution was analyzed by quantifying ultraviolet (UV) absorbance at 220 nm using a Biospec Nano UV spectrometer (Shimadzu).

### 2.2 Radiolabeling of PSMA-617

PSMA-617 was radiolabeled with ^89^Zr in a mixture of HEPES buffer and organic solvent. Solvent choice was determined by the results of the following experiments. One microliter of 10^-2^ mol/L PSMA-617 (10 nmol), 449 μL of 0.5 mol/L HEPES buffer (pH 7.0), 500 μL of organic solvent, and 50 μL of ^89^Zr chloride solution were added to tubes and reacted at 90°C for 30 min. Methanol (MeOH), ethanol (EtOH), N, N-dimethylformamide (DMF), N-methylpyrrolidone (NMP), and dimethylsulfoxide (DMSO) were tested. Water was also tested as a reference. The oxalate concentration was controlled by adding a volume of 10^-2^ mol/L oxalic acid to obtain the following final oxalic acid concentrations: 0 mol/L, 10^-5^ mol/L, 5 × 10^-5^ mol/L, and 10^-6^ mol/L. The radiochemical yield (RCY) was evaluated by instant thin-layer chromatography (ITLC) using ITLC-SG (Agilent) and 1 mol/L ammonium acetate/methanol (1:1) as the mobile phase.

^89^Zr-PSMA-617 for *in vivo* and *in vitro* experiments was prepared as follows. One microliter of 10^-2^ mol/L PSMA-617 (10 nmol), 699 μL of 0.5 mol/L HEPES buffer (pH 7.0), 1,000 μL of DMSO, and 300 μL of ^89^Zr chloride solution were added to tubes and reacted at 90°C for 30 min. Oxalic acid was not added. The purification was performed using high-performance liquid chromatography (HPLC) with 2 mmol/L acetic acid (solvent A) and methanol (solvent B), and a 5–100% solvent B gradient over 20 min at a flow rate of 1 mL/min. The fractionated ^89^Zr-PSMA-617 solution was dried *in vacuo.* Finally, ^89^Zr-PSMA-617 was redissolved by saline and used for the experiments here described.

### 2.3 Cell culture

The PSMA-positive (PSMA+) LNCaP cell line (metastatic lesion of human prostatic adenocarcinoma, ATCC CRL-1740) and the PSMA-negative (PSMA-) PC-3 cell line (bone metastasis of a grade IV prostatic adenocarcinoma, ATCC CRL-1435) were cultured in RPMI-1640 medium supplemented with heat-inactivated 10% fetal calf serum. Cell culture was performed at 37°C in a 5% CO_2_ atmosphere. The cells were harvested using trypsin-ethylenediaminetetraacetic acid (trypsin-EDTA; 0.25% trypsin, 0.02% EDTA). The expression of PSMA protein in each cell line was investigated by western blot using anti-PSMA rabbit antibody (Cell Signaling Technology) [11].

### 2.4 Cell-based assays

The dissociation constant (*K*_d_) and internalization of ^89^Zr-PSMA-617 was determined using cell-based assays. Twenty four-well plates seeded with 10^5^ LNCaP cells were used for both assays. The plates were precoated with poly-L-lysine to improve cell adherence [12]; the cells were then grown overnight at 37°C and 5% CO_2_. Blocking experiments were also conducted for each assay to assess non-specific binding. The medium used in the blocking assay experiments included 2-phosphonomethyl pentanedioic acid (2-PMPA, 200 μmol/L).

The of *K*_d_ ^89^Zr-PSMA-617 was evaluated based on methods reported previously [13].

After removing the supernatant, different concentrations (0.12–500 nmol/L) of ^89^Zr-PSMA-617 in 500 μL RPMI-1640 medium was added. The well plates were incubated for 30 min at 4°C. The supernatants were then removed and the cells were washed twice with ice-cold PBS followed by the addition of NaOH (0.3 mol/L, 600 μL) to each well. The cell suspensions were transferred to microtubes for measurement in a gamma counter (Hidex). The *K*_d_ values were determined by plotting specific-binding (total binding minus non-specific binding) against the molar concentration of the added radioligands followed by nonlinear regression analysis using GraphPad Prism 7 software.

Internalization of ^89^Zr-PSMA-617 to LNCaP cells was evaluated by methods reported previously [14]. After removing the supernatant, 32 nM of ^89^Zr-PSMA-617 in 250 μL Opti-MEM was added, and well plates were incubated for 45 min at 37°C. The cells were washed four times with ice-cold PBS and then washed twice with 50 mM glycine (pH 2.8). After washing the cells with PBS, the internalized fraction was determined by lysis of the LNCaP cells using 0.3 M NaOH. The radioactivity collected from the glycine and hydroxide fractions was measured in a gamma counter. The specific cell surface binding and specific internalization were calculated by subtracting the nonspecific cell surface binding and the nonspecific internalized fraction and expressed as %ID/10^6^ LNCaP cells.

### 2.5 Mouse tumor model

This study was approved by the Animal Experimentation Committee on Isotope Science Center, The University of Tokyo. All animal experiments were carried out according to The University of Tokyo Animal Experimentation Regulations and ARRIVE guidelines.

Seven-week-old male nude mice (BALB/c *nu*/*nu*) were purchased from Japan SLC Inc. Mouse tumor models were established by subcutaneously injection both of LNCaP (5 × 10^6^ cells) to the right shoulder and PC-3 (5 × 10^6^ cells) to left shoulder. Each cell line was suspended in 50% Matrigel (Corning) before injection. This mouse tumor model was used for biodistribution and PET imaging studies of ^89^Zr-PSMA-617.

### 2.6 PET imaging and biodistribution studies

In order to evaluate whether ^89^Zr-PSMA-617 can visualize PSMA positive tumors, preliminary small-animal PET imaging studies were performed. ^89^Zr-PSMA-617 saline solution (~5 nmol per mouse, 3–7 MBq per mouse, N = 5) were injected via a lateral tail vein into mice bearing both LNCaP (right shoulder, PSMA+) and PC-3 (left shoulder, PSMA-) tumor xenografts. The anesthetized animals (2% isoflurane) were promptly placed into the PET scanner (Clairvivo PET, Shimadzu Corporation) to perform a 30 min PET scan at 24 h p.i.

The standardized uptake value (SUV) for PSMA+, PSMA-, and kidney at p.i. 24 h was calculated as SUV_max_, SUV_mean_ (95%), and SUV_peak_. SUV_max_ is the SUV of the hottest voxel within a defined volume of interest (VOI). SUV_mean_ (95%) is the SUV for voxels with a signal higher than 95% of the maximum intensity in VOI. SUV_peak_ is the average SUV computed within a fixed-size VOI containing the hottest pixel value. A sphere around the hottest pixel with a 1.6-mm diameter was set as the fixed-size VOI in this study.

After the PET scans 24 h after injection, the animals were sacrificed and the organs of interest were dissected and weighed. The radioactivity of each organ was measured using a gamma counter (Cobra quantum, PerkinElmer). The mean value and the standard deviation of %ID/g was calculated.

### 2.7 Statistical analysis

All statistical analyses in this study were conducted using GraphPad Prism 7. The standard deviation was adopted as the error range. For the PET imaging experiments, the difference in the accumulation of ^89^Zr-PSMA-617 in PSMA+ and PSMA-tumors was assessed using a paired-*t*-test.

## 3 Results

### 3.1 Production and purification of ^89^Zr

The Ge detector observed only specific γ-ray energies from ^89^Zr (e.g., 511, 909 keV). An average 69 ± 4% of activity was recovered after ^89^Zr purification. The oxalic acid content of the purified ^89^Zr solution was below 10^-5^ mol/L. These data confirm the high radionuclidic purity and low oxalic acid concentration of the purified ^89^Zr solution.

### 3.2 Radiolabeling of PSMA-617

In order to obtain a high radiochemical yield (RCY) of ^89^Zr-PSMA-617, we set out to identify the optimum reaction buffer. As summarized in Figure 1, RCY was significantly increased by using organic solvent. DMSO had no significant effect on RCY, at any of the oxalate concentrations used. The overall RCY of the ^89^Zr-PSMA-617 saline solution, including the HPLC separation process, was 70 ± 9%. Our data show that a high RCY of ^89^Zr-PSMA-617 can be successfully prepared using a mixture of DMSO and HEPES buffers. Quality control of ^89^Zr-PSMA-617 prepared in this study was performed using radio-HPLC. Chromatographs of non-radioactive PSMA-617, free ^89^ZrCl_4_, and ^89^Zr-PSMA-617 are shown in the Supplementary Material (Figure S1). The retention time of ^89^Zr-PSMA-617 was consistent with that of PSMA-617.

**FIGURE 1.**
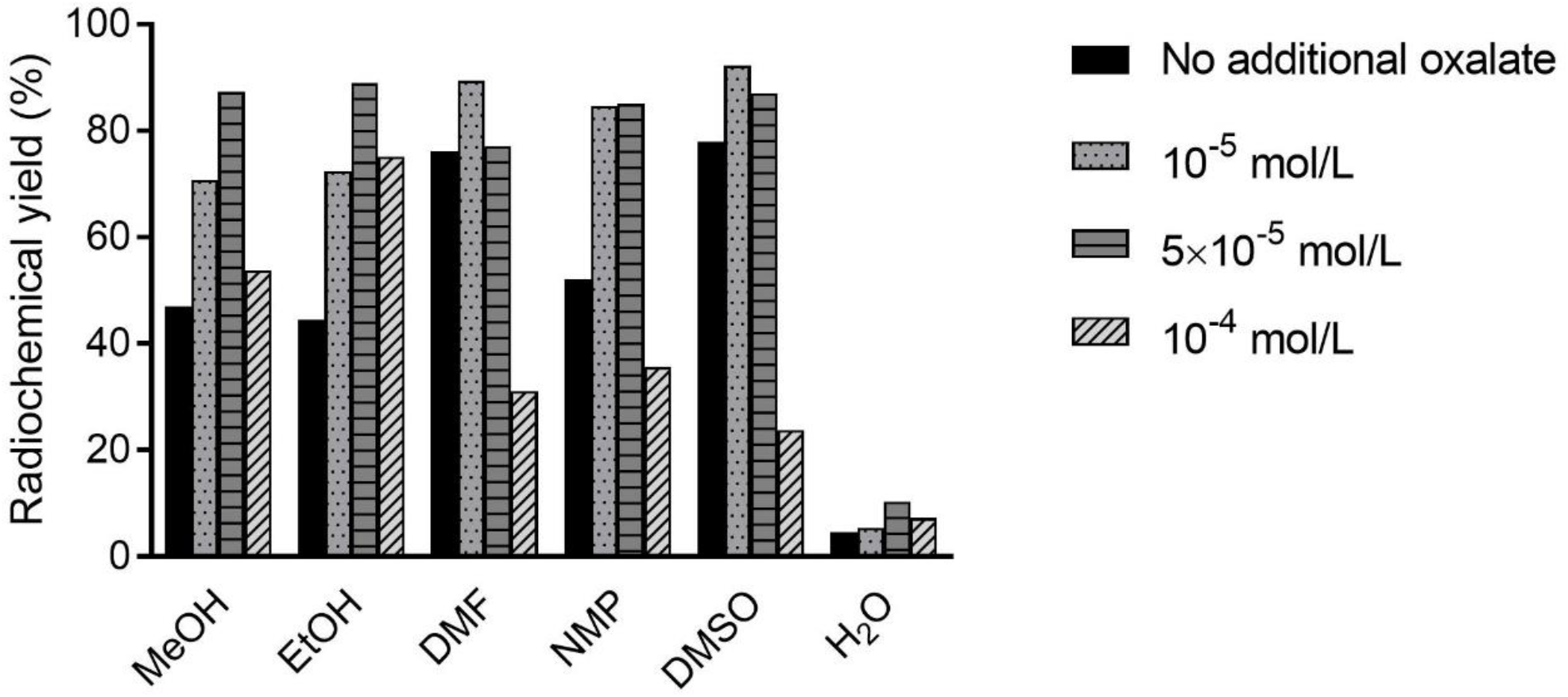
The radiochemical yield of ^89^Zr-PSMA-617 reacted in a mixture of HEPES buffer and organic solvent with different oxalate concentrations.

### 3.3 Cell binding assay

A cell binding assay was performed to determine the *K*_d_ of ^89^Zr-PSMA-617. Western blot experiments confirmed the expression of PSMA protein in LNCaP cells and its absence in PC-3 cells (Figure S2). The binding assay revealed a *K*_d_ of 6.8 nM (95% confidence interval: 2.6–17 nM) for ^89^Zr-PSMA-617 (Figure 2).

**FIGURE 2.**
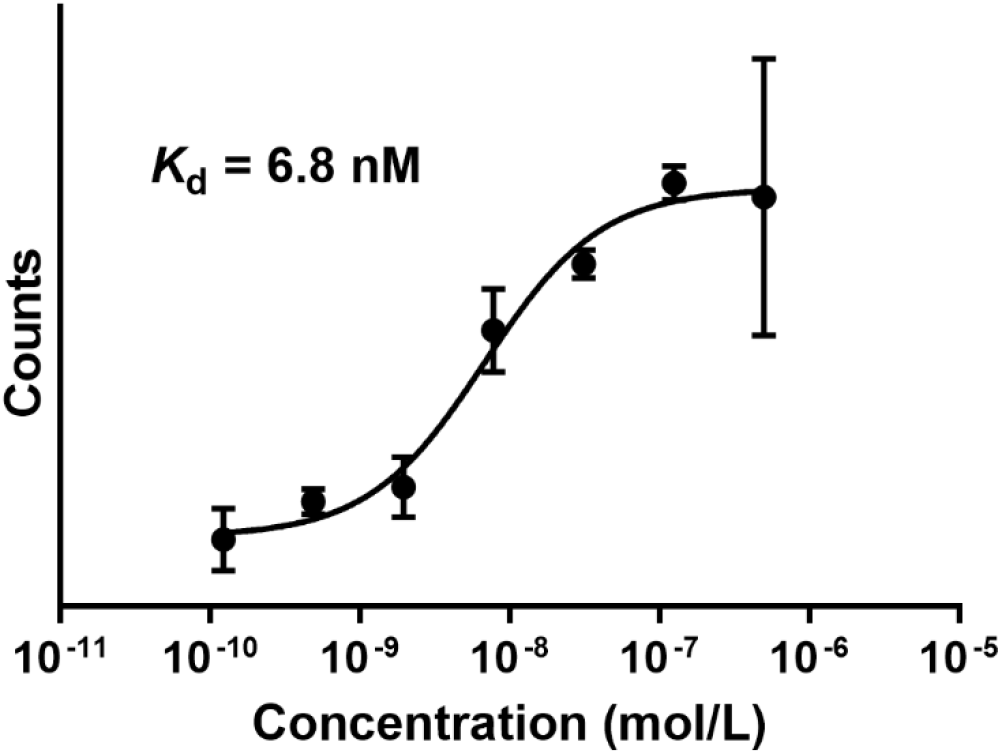
A representative graph of experiments performed to determine the *K*_d_ values showing counts measured in cell samples exposed to different concentrations of ^89^Zr-PSMA-617. N=3 for both experiments and error bars indicate the standard deviation.

In LNCaP, the specific cell surface binding of ^89^Zr-PSMA-617 was 19.17 ± 0.65%ID/10^6^ cells, and the specific internalization was 12.66 ± 0.60%/10^6^ cells.

### 3.4 PET imaging and biodistribution studies

Delayed-PET imaging experiments using LNCaP (PSMA+) and PC-3 (PSMA-) tumor-bearing mice were performed 24 h after injection. Figure 3 (a) shows an example of maximum intensity projection (MIP) PET/CT images; these images were obtained using the mouse shown in Figure 3 (b). Significant accumulation of activity was found in LNCaP tumors (PSMA+) and kidneys but not in PC-3 tumors (PSMA-). The SUVs of other tissues were all below 0.1. The panels in Figure 3 (c–e) summarize the SUVs for PSMA+ cells, PSMA-cells, and kidney at 24 h p.i. There was a significant difference in SUV between PSMA+ and PSMA-samples (P < 0.01).

**FIGURE 3.**
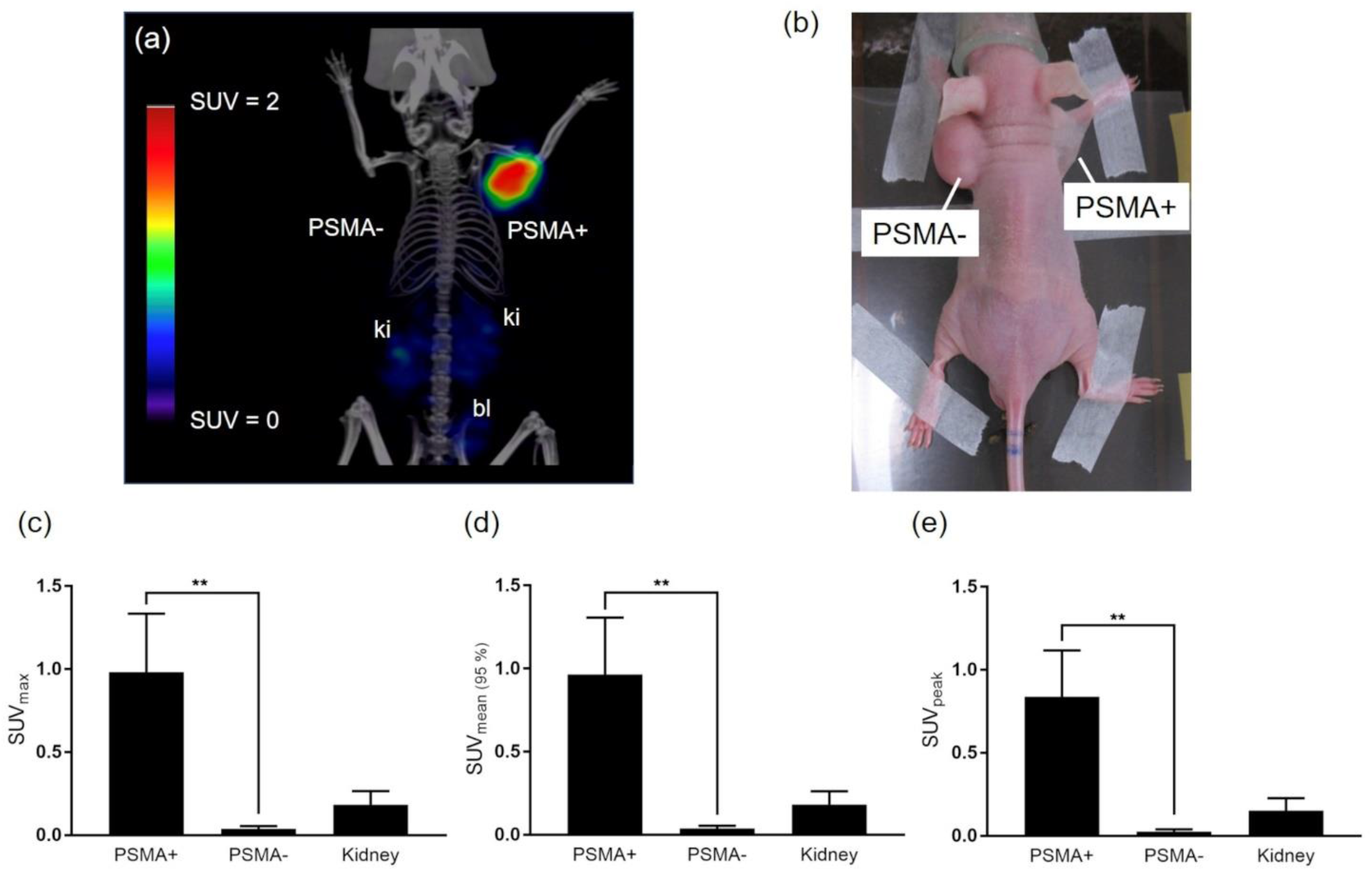
PET/CT image of ^89^Zr-PSMA-617 at 24 h and comparisons of SUV in tumors and kidney. (a) PET/CT image of LNCaP (PSMA+) and PC-3 (PSMA-) tumor-bearing mice at 24 h p.i. (Abbreviations: bl = bladder, ki = kidney.) (b) A picture of the mouse used for PET/CT imaging experiments. (c–e) Comparison of SUV for PSMA+, PSMA-, and kidney at 24 h p.i. N = 5; error bars indicate the standard deviation. SUV was calculated as SUV_max_, SUV_mean_ (95%), and SUV_peak_. P-values lower than 0.01 are represented by two asterisks (**).

Phantom experiments using NEMA PET Small Animal Phantom imaging device are shown in the Supplementary material (Figure S3).

The distribution profiles of ^89^Zr-PSMA-617 in LNCaP (PSMA+) and PC-3 (PSMA-) tumor-bearing mice 24 h after injection are shown in Figure 4. The highest accumulation (1.76 ± 0.61%ID/g) was in LNCaP tumors (PSMA+) and the second highest (0.46 ± 0.15%ID/g) was in the kidneys. The accumulation in PC-3 tumors (PSMA-) and other tissues was similar to the background level (below 0.1%ID/g).

**FIGURE 4.**
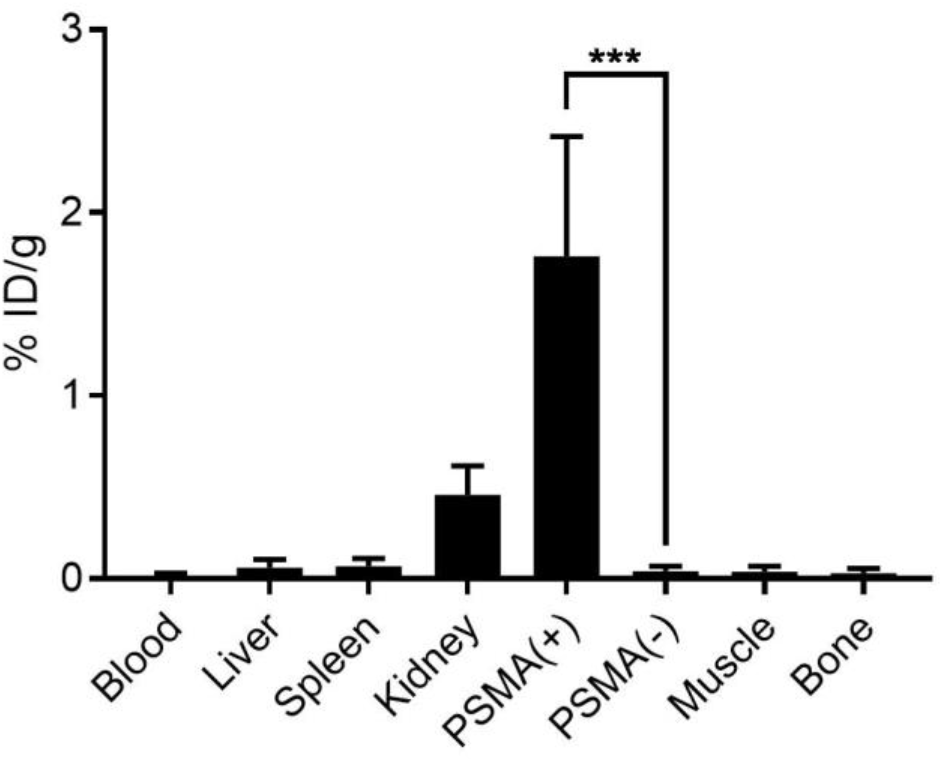
Biodistribution of ^89^Zr-PSMA-617 in LNCaP-(PSMA+) and PC-3 (PSMA-) tumor-bearing mice at 24. h p.i. N = 7; error bars indicate the standard deviation. P-values lower than 0.001 are represented by three asterisks (***).

## 4 Discussion

Here, we have established a method of ^89^Zr-PSMA-617 synthesis using an organic solvent. In previous studies, radiolabeling of the DOTA chelator with ^89^Zr was performed in the presence of HEPES buffer alone [15]. Under this condition, very high concentrations of DOTA (higher than 10^-4^ mol/L (100 nmol/mL)) were required to obtain a high radiochemical yield (RCY) (> 90%) [16]. Considering that a small mass of PSMA-617 or PSMA-11 (typically 2 nmol) is administered to each patient, such a high DOTA concentration results in low specific radioactivity (SA) [7,17,18]. In contrast, we demonstrate in the current report that ^89^Zr radiolabeling of PSMA-617 can be achieved with a DOTA concentration of 5 × 10^-6^ mol/L (5 nmol/mL), which is one-twentieth of the conventional concentration, by using DMSO. Our improved technique effectively reduces the PSMA-617 concentration required for efficient ^89^Zr radiolabeling to the concentration commonly used for ^68^Ga radiolabeling (5–50 nmol/mL) [7,14].

We speculate that the irreversible formation of ^89^Zr hydroxide ^89^Zr(OH)4 was responsible for the low RCY when organic solvent is not used. Due to the poor aqueous solubility of Zr hydroxide [19], ^89^Zr would be poorly reactive with DOTA once ^89^Zr hydroxide is formed. The ability to achieve high RCYs when an organic solvent is used is probably due to inhibition of ^89^Zr hydroxide formation. Since Zr oxalate is more readily formed than Zr hydroxide [20], a slight amount of oxalate might improve RCY. Indeed, RCY was consistently high when using DMSO, irrespective of the oxalate concentrations tested in this study (Figure 1). DMSO is a solvent frequently used in pharmaceutical synthesis. Although the DMSO concentration in the final formulation needs to be controlled, we suggest that radiolabeling using DMSO would be acceptable in future clinical studies.

The *K*_d_ values of ^89^Zr-PSMA-617 (6.8 nM) agreed with those of PSMA-617 labeled with other radionuclides [13,21–23]. The specific internalization of ^89^Zr-PSMA-617 (12.66 ± 0.60%/10^6^ cells) was close to ^177^Lu-PSMA-617 (16.17 ± 3.66%/10^6^ cells). While DOTA-conjugated peptides are usually combined with divalent or trivalent metal ions, ^89^Zr^4+^ (tetravalent) labeled DOTA-conjugated peptides also exhibited comparable cell binding ability and internalization.

As we anticipated, PSMA+ lesions could be visualized using delayed-PET imaging with long-lived ^89^Zr-PSMA-617 while unfavorable accumulation in the bladder was avoided (Figure 4). In contrast, clinical studies show that radioactive ^68^Ga- and ^44^Sc-PSMA-617 remain in the bladder at the time of PET-scanning [7,24]. Therefore, we propose that ^89^Zr-PSMA-617 is a more suitable reagent for the prediction of cytotoxic doses of PSMA-RLT. However, we note that the PET imaging studies of ^89^Zr-PSMA-617 were performed in animal experiments only, and future clinical studies will be required to validate our hypothesis. In addition, the SA of ^89^Zr-PSMA-617 prepared in this study was relatively low (~ 1 GBq/μmol) because ^89^Zr purification was not automated and we were limited in the amount of radioactivity that could be handled due to the radiation safety guidelines of our facility. Thus, a relatively large amount of radioligand (~5 nmol per mouse) was administrated. The accumulation of ^89^Zr-PSMA-617 (1.76 ± 0.61%ID/g) in tumors was lower than that of ^177^Lu-PSMA-617 (10.58 ± 4.50%ID/g) [7]. This discrepancy could be attributed to differences in the number of injected radioligands (^177^Lu-PSMA-617: 0.06 nmol). Research interest in future studies will be the automation of ^89^Zr purification to the chloride form and comparisons of ^89^Zr-PSMA-617 and ^177^Lu-PSMA-617 biodistribution when lower levels are administered (> 1 nmol per mouse).

Delayed-PET imaging at 24 h p.i. have been achieved by ^64^Cu-PSMA-617. However, the biodistribution profile of ^64^Cu-PSMA-617 at 24 h p.i. was different to that of ^89^Zr-PSMA-617. Accumulation of ^89^Zr-PSMA-617 in the liver was negligible (below 0.1%ID/g), while that of ^64^Cu-PSMA-617 at 24 h p.i. was considerably higher (9.08%ID/g) [21]. The increased liver uptake of ^64^Cu-PSMA-617 is most probably due to free radionuclides, because ^64^Cu-Macrocyclic complexes, which have limited in vivo stability, are dissociated by superoxide dismutase in the liver [25,26]. In contrast, in vivo dissociation of ^89^Zr-PSMA-617 was inferred to be low. Free ^89^Zr in plasma accumulates in the bone and liver [27,28]. Our biodistribution experiment revealed negligible accumulation (below 0.1%ID/g) in these organs, which also highlights the biochemical stability of ^89^Zr-PSMA-617 *in vivo*. The low accumulation in bone and liver agrees well with the amount of ^89^Zr-DOTA complex in these organs at 24 h p.i. (bone: 0.036%ID/g, liver: 0.068%ID/g) [15].

Radioactive urine in the bladder can also complicate the evaluation of signals coming from adjacent organs, such as the prostate. Additionally, urine spots in the ureters could be misdiagnosed as lymph node metastases in human PET images of PSMA. These issues cannot be avoided by using PSMA ligands labeled with ^18^F or ^68^Ga due to their short half-lives. In contrast, ^89^Zr-PSMA-617 allows the visualization of PSMA expression in primary prostate lesions, because the long half-life of ^89^Zr allows scanning to be performed after all radioactive urine has been completely excreted. Thus, the use of ^89^Zr-PSMA-617 could be extended to the diagnostics of primary lesions. However, it should be kept in mind that PMSA expression is apparently high in patients with benign prostatic hyperplasia (BPH) [29]. Future human studies with ^89^Zr-PSMA-617 are required to clarify its utility in this context.

## 5 Conclusion

Herein, we have described the preparation and use of ^89^Zr-PSMA-617 for PET imaging studies. The preparation of ^89^Zr-PSMA-617 was achieved in this study for the first time by using the mixture solvent of HEPES buffer and DMSO. ^89^Zr-PSMA-617 prepared in this study clearly visualize the PSMA positive LNCaP tumors without accumulation in bladder. The relatively long half-life of ^89^Zr circumvents the problem of unfavorable bladder accumulation, and ^89^Zr-PSMA-617 could therefore be an optimum choice for pre-therapeutic dosimetry of ^177^Lu-PSMA-617. For the same reason, ^89^Zr-PSMA-617 could be used to visualize primary prostate lesions in PC patients who are being evaluated for the presence of metastases. The use of ^89^Zr-PSMA-617 could also be extended to the diagnostics of primary lesions.

## Disclosure

Ryota Imura and Hiroyuki Ida are employees of JFE Engineering Corporation. This research was partially conducted with research funds of the JFE Engineering Corporation.

## Acknowledgments

This research was conducted with research funds from JFE Engineering Corporation, which employs two of this paper’s authors (Ryota Imura and Hiroyuki Ida).

We gratefully thank Professor Hiroshi Watabe (Cyclotron and Radioisotope Center, Tohoku University) for supporting the operation of the PET scanner and data analysis. We also thank Toshifumi Omura and Toru Matsumoto (JFE Engineering Corporation) for technical help.

## Visual Abstract

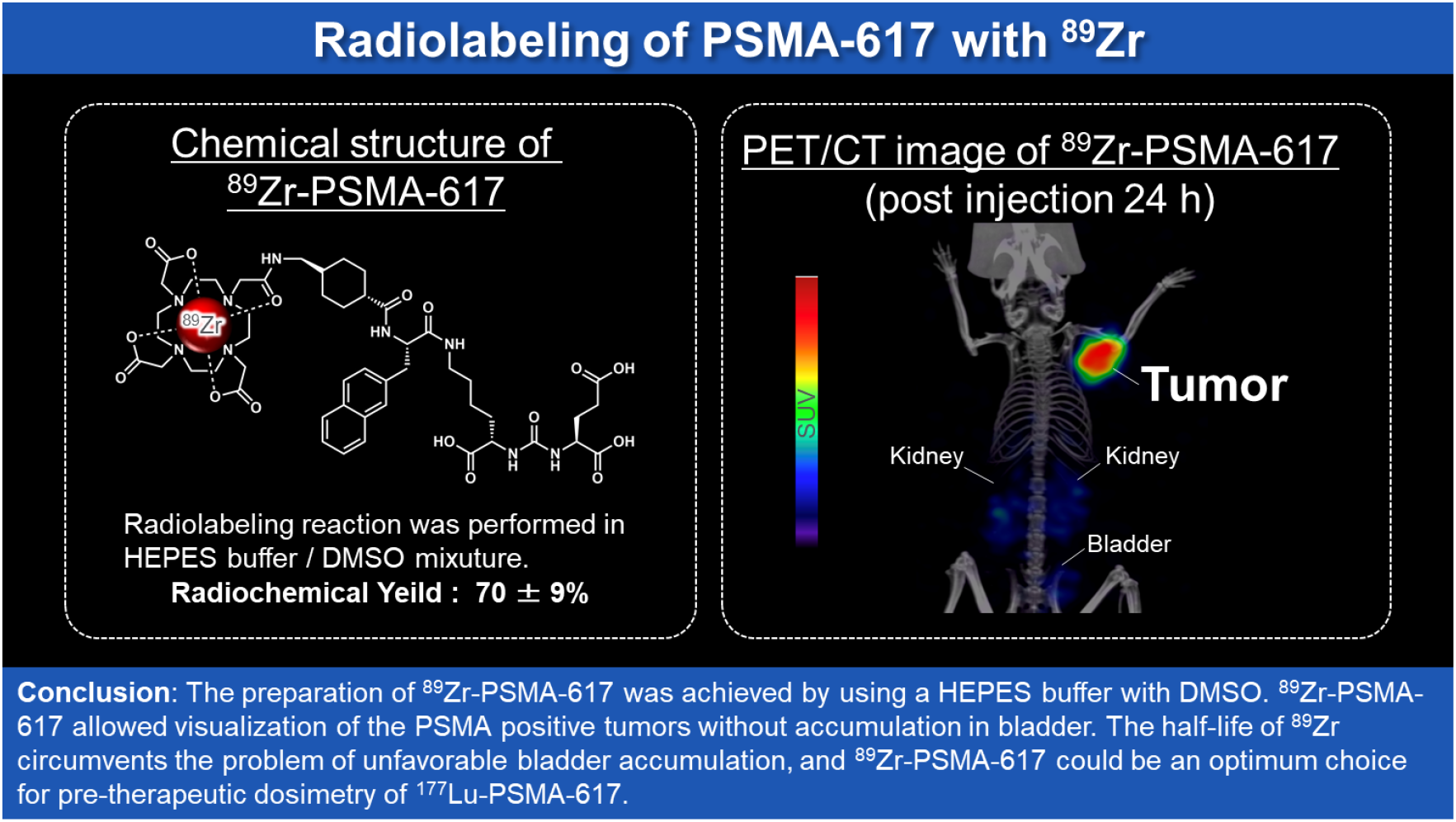

## Notes

**Funding** This research was conducted with research funds from JFE Engineering Corporation. No other potential conflicts of interest relevant to this article exist.

## References

[1] Silver DA, Pellicer I, Fair WR, Heston WD, Cordon-Cardo C. Prostate-specific membrane antigen expression in normal and malignant human tissues. Clin Cancer Res 1997;3:81–5.

[2] Afshar-Oromieh A, Hetzheim H, Kratochwil C, Benesova M, Eder M, Neels OC, et al. The Theranostic PSMA Ligand PSMA-617 in the Diagnosis of Prostate Cancer by PET/CT: Biodistribution in Humans, Radiation Dosimetry, and First Evaluation of Tumor Lesions. J Nucl Med 2015;56:1697–705. https://doi.org/10.2967/jnumed.115.161299.

[3] O’Keefe DS, Bacich DJ, Heston WDW. Comparative analysis of prostate-specific membrane antigen (PSMA) versus a prostate-specific membrane antigen-like gene. Prostate 2004;58:200–10. https://doi.org/10.1002/pros.10319.

[4] Foss CA, Mease RC, Fan H, Wang Y, Ravert HT, Dannals RF, et al. Radiolabeled Small-Molecule Ligands for Prostate-Specific Membrane Antigen: In vivo Imaging in Experimental Models of Prostate Cancer. Clin Cancer Res 2005;11:4022–8. https://doi.org/10.1158/1078-0432.ccr-04-2690.

[5] Hillier SM, Maresca KP, Femia FJ, Marquis JC, Foss CA, Nguyen N, et al. Preclinical Evaluation of Novel Glutamate-Urea-Lysine Analogues That Target Prostate-Specific Membrane Antigen as Molecular Imaging Pharmaceuticals for Prostate Cancer. Cancer Res 2009;69:6932–40. https://doi.org/10.1158/0008-5472.can-09-1682.

[6] Banerjee SR, Pullambhatla M, Byun Y, Nimmagadda S, Green G, Fox JJ, et al. 68Ga-Labeled Inhibitors of Prostate-Specific Membrane Antigen (PSMA) for Imaging Prostate Cancer. J Med Chem 2010;53:5333–41. https://doi.org/10.1021/jm100623e.

[7] Benešová M, Schäfer M, Bauder-Wüst U, Afshar-Oromieh A, Kratochwil C, Mier W, et al. Preclinical Evaluation of a Tailor-Made DOTA-Conjugated PSMA Inhibitor with Optimized Linker Moiety for Imaging and Endoradiotherapy of Prostate Cancer. J Nucl Med 2015;56:914–20. https://doi.org/10.2967/jnumed.114.147413.

[8] Ahmadzadehfar H, Rahbar K, Kürpig S, Bögemann M, Claesener M, Eppard E, et al. Early side effects and first results of radioligand therapy with 177Lu-DKFZ-617 PSMA of castrate-resistant metastatic prostate cancer: a two-centre study. EJNMMI Res 2015;5:36. https://doi.org/10.1186/s13550-015-0114-2.

[9] Disselhorst JA, Brom M, Laverman P, Slump CH, Boerman OC, Oyen WJG, et al. Image-Quality Assessment for Several Positron Emitters Using the NEMA NU 4-2008 Standards in the Siemens Inveon Small-Animal PET Scanner. J Nucl Med 2010;51:610–7. https://doi.org/10.2967/jnumed.109.068858.

[10] Holland JP, Sheh Y, Lewis JS. Standardized methods for the production of high specific-activity zirconium-89. Nucl Med Biol 2009;36:729–39. https://doi.org/10.1016/j.nucmedbio.2009.05.007.

[11] Yamada T, Imamachi N, Imamura K, Taniue K, Kawamura T, Suzuki Y, et al. Systematic Analysis of Targets of Pumilio-Mediated mRNA Decay Reveals that PUM1 Repression by DNA Damage Activates Translesion Synthesis. Cell Reports 2020;31:107542. https://doi.org/10.1016/j.celrep.2020.107542.

[12] Liberio MS, Sadowski MC, Soekmadji C, Davis RA, Nelson CC. Differential Effects of Tissue Culture Coating Substrates on Prostate Cancer Cell Adherence, Morphology and Behavior. PLoS One 2014;9:e112122. https://doi.org/10.1371/journal.pone.0112122.

[13] Umbricht CA, Benešová M, Schmid RM, Türler A, Schibli R, Meulen NP van der, et al. 44Sc-PSMA-617 for radiotheragnostics in tandem with 177Lu-PSMA-617— preclinical investigations in comparison with 68Ga-PSMA-11 and 68Ga-PSMA-617. EJNMMI Res 2017;7:9. https://doi.org/10.1186/s13550-017-0257-4.

[14] Benešová M, Bauder-Wüst U, Schäfer M, Klika KD, Mier W, Haberkorn U, et al. Linker Modification Strategies To Control the Prostate-Specific Membrane Antigen (PSMA)-Targeting and Pharmacokinetic Properties of DOTA-Conjugated PSMA Inhibitors. J Med Chem 2016;59:1761–75. https://doi.org/10.1021/acs.jmedchem.5b01210.

[15] Pandya DN, Bhatt N, Yuan H, Day CS, Ehrmann BM, Wright M, et al. Zirconium tetraazamacrocycle complexes display extraordinary stability and provide a new strategy for zirconium-89-based radiopharmaceutical development. Chem Sci 2016;8:2309–14. https://doi.org/10.1039/c6sc04128k.

[16] Graves SA, Kutyreff C, Barrett KE, Hernandez R, Ellison PA, Happel S, et al. Evaluation of a chloride-based 89Zr isolation strategy using a tributyl phosphate (TBP)-functionalized extraction resin. Nucl Med Biol 2018. https://doi.org/10.1016/j.nucmedbio.2018.06.003.

[17] Afshar-Oromieh A, Zechmann CM, Malcher A, Eder M, Eisenhut M, Linhart HG, et al. Comparison of PET imaging with a 68Ga-labelled PSMA ligand and 18F-choline-based PET/CT for the diagnosis of recurrent prostate cancer. Eur J Nucl Med Mol Imaging 2014;41:11–20. https://doi.org/10.1007/s00259-013-2525-5.

[18] Afshar-Oromieh A, Avtzi E, Giesel FL, Holland-Letz T, Linhart HG, Eder M, et al. The diagnostic value of PET/CT imaging with the 68Ga-labelled PSMA ligand HBED-CC in the diagnosis of recurrent prostate cancer. Eur J Nucl Med Mol Imaging 2015;42:197–209. https://doi.org/10.1007/s00259-014-2949-6.

[19] Sasaki T, Kobayashi T, Takagi I, Moriyama H. Solubility measurement of zirconium(IV) hydrous oxide. Radiochim Acta 2006;94:489–94. https://doi.org/10.1524/ract.2006.94.9-11.489.

[20] Koyashi T, Sasaki T, Takagi I, Moriyama H. Zirconium Solubility in Ternary Aqueous System of Zr(IV)-OH-Carboxylates. J Nucl Sci Technol 2009;46:142–8. https://doi.org/10.1080/18811248.2007.9711515.

[21] Cui C, Hanyu M, Hatori A, Zhang Y, Xie L, Ohya T, et al. Synthesis and evaluation of [64Cu]PSMA-617 targeted for prostate-specific membrane antigen in prostate cancer. Am J Nucl Medicine Mol Imaging 2017;7:40–52.

[22] Tönnesmann R, Meyer PT, Eder M, Baranski A-C. [177Lu]Lu-PSMA-617 Salivary Gland Uptake Characterized by Quantitative In Vitro Autoradiography. Pharm 2019;12:18. https://doi.org/10.3390/ph12010018.

[23] Gourni E, Canovas C, Goncalves V, Denat F, Meyer PT, Maecke HR. (R)-NODAGA-PSMA: A Versatile Precursor for Radiometal Labeling and Nuclear Imaging of PSMA-Positive Tumors. PLoS One 2015;10:e0145755. https://doi.org/10.1371/journal.pone.0145755.

[24] Eppard E, Fuente A de la, Benešová M, Khawar A, Bundschuh RA, Gärtner FC, et al. Clinical Translation and First In-Human Use of [44Sc]Sc-PSMA-617 for PET Imaging of Metastasized Castrate-Resistant Prostate Cancer. Theranostics 2017;7:4359–69. https://doi.org/10.7150/thno.20586.

[25] Bass LA, Wang M, Welch MJ, Anderson CJ. In Vivo Transchelation of Copper-64 from TETA-Octreotide to Superoxide Dismutase in Rat Liver. Bioconjugate Chem 2000;11:527–32. https://doi.org/10.1021/bc990167l.

[26] Watabe T, Liu Y, Kaneda-Nakashima K, Shirakami Y, Lindner T, Ooe K, et al. Theranostics targeting fibroblast activation protein in the tumor stroma: 64Cu and 225Ac labelled FAPI-04 in pancreatic cancer xenograft mouse models. J Nucl Med 2019:jnumed.119.233122. https://doi.org/10.2967/jnumed.119.233122.

[27] Abou DS, Ku T, Smith-Jones PM. In vivo biodistribution and accumulation of 89Zr in mice. Nucl Med Biol 2011;38:675–81. https://doi.org/10.1016/j.nucmedbio.2010.12.011.

[28] Holland JP, Divilov V, Bander NH, Smith-Jones PM, Larson SM, Lewis JS. 89Zr-DFO-J591 for ImmunoPET of Prostate-Specific Membrane Antigen Expression In Vivo. J Nucl Med 2010;51:1293–300. https://doi.org/10.2967/jnumed.110.076174.

[29] Rowe SP, Gage KL, Faraj SF, Macura KJ, Cornish TC, Gonzalez-Roibon N, et al. 18F-DCFBC PET/CT for PSMA-Based Detection and Characterization of Primary Prostate Cancer. J Nucl Med 2015;56:1003–10. https://doi.org/10.2967/jnumed.115.154336.

