## supplemental material for "Radiolabeling of PSMA-617 with ^89^Zr: A Novel Use of DMSO for Radiochemical Yield Enhancement and Preliminary Small-Animal PET Results"

### Supplementary Material

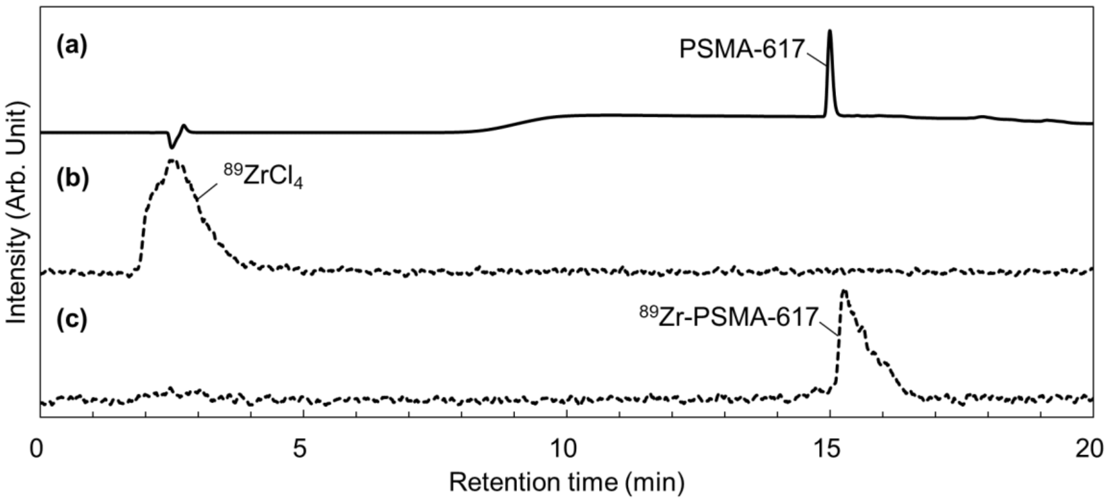


FIGURE S1. Quality control of $^{89}$Zr-PSMA-617 prepared by the methods described in the main article. The measurements were performed using radio-HPLC with water containing 0.1% tetrafluoroacetic acid (solvent A) and acetonitrile containing tetrafluoroacetic acid (solvent B), and a 5–90% solvent B gradient over 20 min at a flow rate of 1 mL/min. The injection volume was 10 µL each. HPLC chromatogram of non-radioactive PSMA-617 monitored by 201 nm UV (a), $^{89}ZrCl_4$ and $^{89}$Zr-PSMA-617 monitored with a radioactivity sensor (b and c, respectively). The retention time of $^{89}$Zr-PSMA-617 was consistent with that of PSMA-617.

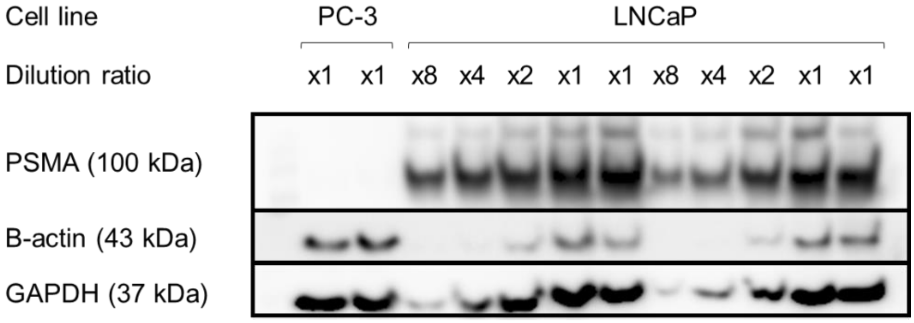


FIGURE S2. The expression of PSMA protein in LNCaP and PC-3 cells was investigated by western blotting. PSMA is in the top image, and control proteins (β-actin and GAPDH) are also indicated. All LNCaP and PC-3 cell samples were derived from the same clones used in the *in vivo* and *in vitro* experiments for $^{89}$Zr-PSMA-617. For LNCaP, 2-fold, 4-fold, and 8-fold diluted samples were prepared. Note the high PSMA expression in LNCaP cells and no expression in PC-3 cells.

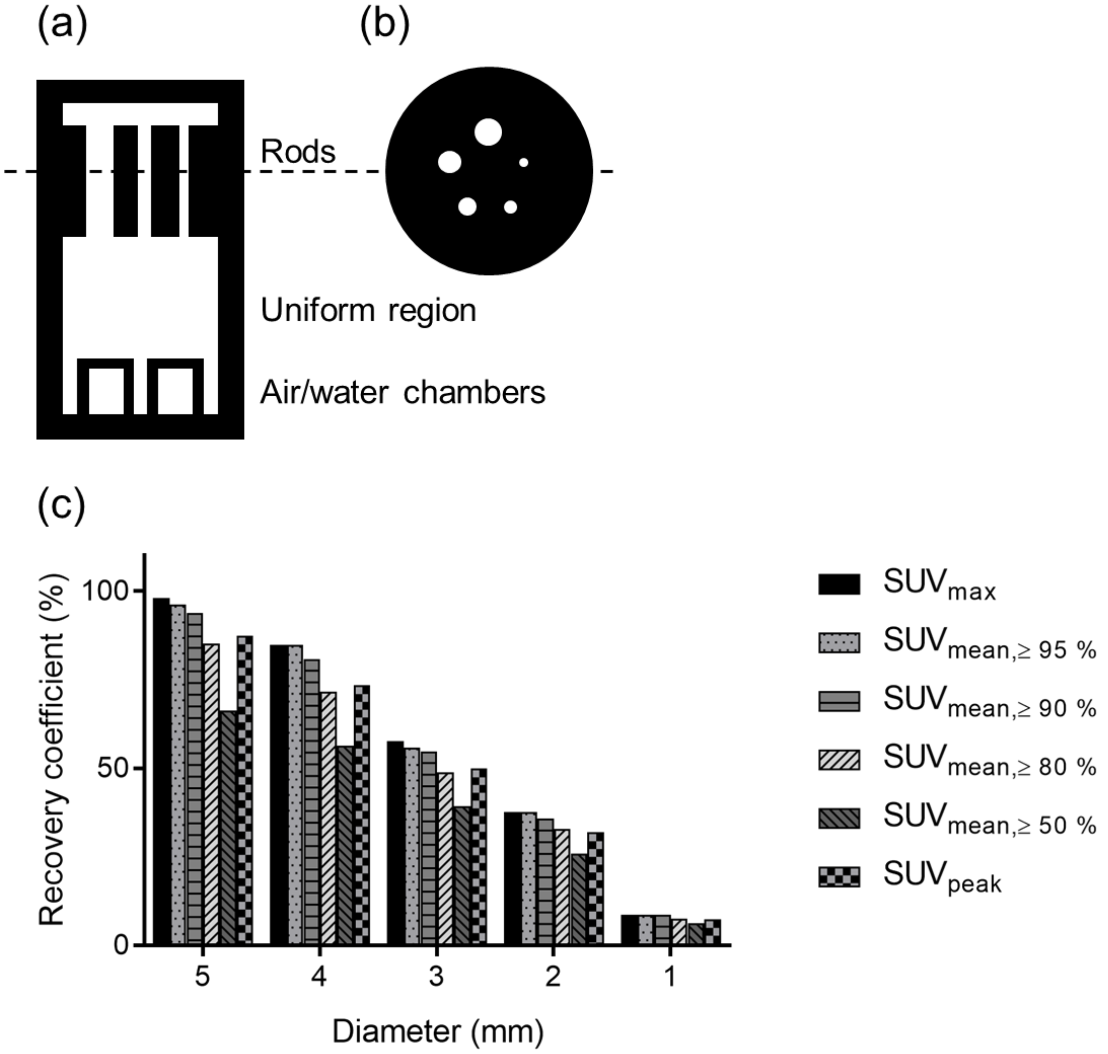


FIGURE S3. The structure of the NEMA NU 4 IQ phantom and the results of the phantom test. Figure S2 (a) and (b) show the structure of the NEMA NU 4 IQ phantom. It contains the following 3 regions: a region with 5 rods (diameters of 1, 2, 3, 4, and 5 mm) to measure recovery coefficients (a; top), a uniform region to measure uniformity (a; middle), and a 2-chamber region (air- and water-filled) to measure background in the reconstructed image (a; bottom). (b) Five-rod region. (c) Recovery coefficients (RCs) evaluated in each of the fillable rods with 1, 2, 3, 4, and 5 mm diameter and 20 mm length. The RC was defined as the ratio between the measured activity concentration in the rods and the activity concentration measured in the uniform area, and is shown for each rod diameter. The ideal RC value is 100%. Volumes of interest (VOIs) were set to the same size as the actual rod dimensions in this test. $SUV_{max}$ was calculated from the pixel with the maximum signal in the VOIs. $SUV_{mean}$ was calculated from pixels with a signal greater than the threshold in the VOIs. For example, $SUV_{mean \geq 95\%}$ was calculated from pixels that had a signal greater than 95% of the maximum signal. $SUV_{peak}$ was calculated as the mean value in pixels within fixed-size VOIs, set as a 1.6 mm-diameter sphere centered on the pixel with the maximum value.
